# *nanotatoR*: A tool for enhanced annotation of genomic structural variants

**DOI:** 10.1101/2020.08.18.254680

**Authors:** Surajit Bhattacharya, Hayk Barseghyan, Emmanuèle C. Délot, Eric Vilain

## Abstract

Whole genome sequencing is effective at identification of small variants but, because it is based on short reads, assessment of structural variants (SVs) is limited. The advent of Optical Genome Mapping (OGM), which utilizes long fluorescently labeled DNA molecules for de novo genome assembly and SV calling, has allowed for increased sensitivity and specificity in SV detection. However, compared to small variant annotation tools, OGM-based SV annotation software has seen little development, and currently available SV annotation tools do not provide sufficient information for determination of variant pathogenicity.

We developed an R-based package, *nanotatoR*, which provides comprehensive annotation as a tool for SV classification. nanotatoR uses both external (DGV; DECIPHER; Bionano Genomics BNDB) and internal (user-defined) databases to estimate SV frequency. Human genome reference GRCh37/38-based BED files are used to annotate SVs with overlapping, upstream, and downstream genes. Overlap percentages and distances for nearest genes are calculated and can be used for filtration. A primary gene list is extracted from public databases based on the patient’s phenotype and used to filter genes overlapping SVs, providing the analyst with an easy way to prioritize variants. If available, expression of overlapping or nearby genes of interest is extracted (e.g. from an RNA-Seq dataset, allowing the user to assess the effects of SVs on the transcriptome). Most quality-control filtration parameters are customizable by the user. The output is given in an Excel file format, subdivided into multiple sheets based on SV type and inheritance pattern (INDELs, inversions, translocations, de novo, etc.).

*nanotatoR* passed all quality and run time criteria of Bioconductor, where it was accepted in the April 2019 release. We evaluated *nanotatoR’s* annotation capabilities using publicly available reference datasets: the singleton sample NA12878, mapped with two types of enzyme labeling, and the NA24143 trio. *nanotatoR* was also able to accurately filter the known pathogenic variants in a cohort of patients with Duchenne Muscular Dystrophy for which we had previously demonstrated the diagnostic ability of OGM. The extensive annotation enables users to rapidly identify potential pathogenic SVs, a critical step toward use of OGM in the clinical setting.

## Introduction

With the advent of the high throughput short-read sequencing (SRS) techniques, identification of molecular underpinnings of genetic disorders has become faster, more accurate and cost-effective [1]. SRS platforms used for whole exome (WES) or genome (WGS) DNA sequencing, produce billion of reads per run, typically limited in length to 100-150 base pairs (bp) [2]. When WES started being used in clinical diagnostic practice, it was reported to be effective in identifying pathogenic genetic variants in approximately 30% of cases [3–5]. Even with technological evolution and more widespread practice, reports of diagnostic yields between 8 and 70%, depending on the disease [6] suggest that a large fraction of cases remain undiagnosed. WGS was shown to be more effective than WES in identifying single nucleotide variants (SNVs) or small insertions and deletions (INDELs) than WES [7, 8]. However, both WES and WGS are ineffective in identification of structural variants (SVs, >50 bps) or copy number variants (CNVs) because short reads cannot span repetitive elements or provide contextual information. Many algorithms have been designed for detection of SVs in short-read-based sequences (69 of them were compared in [9]). However, performance analyses highlight their limitations such as low concordance, poor precision, and high rate of false positive calls [9, 10]. Another benchmarking study comparing 10 different SV callers against robust truth sets showed that the total number of calls made by the different algorithms varied by greater than two orders of magnitude [11]. Region of genome analyzed (repeats vs. high-complexity regions), noise of data (platform-specific sequencing or assembly errors), complexity of the SV, and library properties (e.g. insert size) all affect specificity, sensitivity and/or processing speed of the various variant-calling algorithms [10].

Chromosomal microarray (CMA) is the established method for high-accuracy detection of CNVs, but it can only identify gains or losses of genetic material and is virtually blind towards identification of balanced rearrangements such as inversions or translocations. CMA clinical application is typically limited to CNVs above 25-50 kb, although higher resolution CNV maps have been built and are being used to design disease-specific paths to diagnostic detection of smaller variants (*e*.*g*. [12]). Breakpoint resolution is limited by the density of probes on the array.

Novel approaches which analyze single, long DNA molecules hold the promise of detecting the previously inaccessible SVs. Long-read sequencing (LRS) technologies such as nanopore-based sequencing (Oxford Nanopore Technologies) or single-molecule real-time sequencing (Pacific Biosciences) have the potential to both detect complex genomic rearrangements and increase SV break point resolution [13, 14]. They have been critical to shed light on the “dark” regions of the genome where short reads had been insufficient for accurate assembly [15]. However, as LRS still isn’t in wide use, SV detection pipelines have seen slower development than SRS-based algorithms, and both quantity and quality of identified SVs vary significantly between tools [16].

In parallel, a method not based on sequencing, optical genome mapping (OGM, Bionano Genomics), provides much higher sensitivity and specificity for identification of large SVs, including balanced events, compared to karyotype, CMA, LRS and SRS [17–20]. For example, a comparison of OGM, PacBio LRS and Illumina-based SRS on the same genome showed that about a third of deletions and three quarters of insertions above 10 kb were detected only by OGM [17]. For OGM, purified high-molecular-weight DNA is fluorescently labeled at specific sequence motifs throughout the genome (reviewed in [21]). The labeled DNA is imaged through nanochannel arrays for *de novo* genome assembly. Assembly and variant calling are performed using algorithms provided by Bionano Genomics and/or tools developed by the community, such as OMblast [22] or OMtools [23]. OGM has been effective in identifying pathogenic variants in patients with cancer [24–26], Duchenne muscular dystrophy [27], and facioscapulohumeral muscular dystrophy [28, 29]. Importantly OGM has allowed refinement of intractable, low-complexity regions of the genome and discovery of genomic content missing in the reference genome assembly [19].

Although, OGM is effective in identifying clinically relevant SVs, the currently available SV annotation tools do not provide sufficient variant information for determination of variant pathogenicity. Here, we report the development of an annotation tool in R language, *nanotatoR*, that provides extensive annotation for SVs identified by OGM. It determines population variant frequency using publicly available databases, as well as user-created internal databases. It offers multiple filtration options based on quality parameters thresholds. It also determines the percentage of overlap of genes with the SV, as well as distance between nearest genes and SV breakpoints, both upstream and downstream. It offers an option for incorporating RNA-Seq read counts, which has been shown to enhance variant classification [6], as well as user-specified disease-specific gene lists extracted from NCBI databases. The final output is provided in an Excel worksheet, with segregated SV types and inheritance patterns, facilitating filtration and identification of pathogenic variants.

## Materials and Methods

### Sample data sets

Optically mapped genomes for 8 different reference human samples were used to construct the internal cohort database for evaluating *nanotatoR’s performance*. All sample datasets, including “Utah woman” (Genome in a Bottle Consortium sample NA12878), “Ashkenazi family” (NA24143 [or GM24143]: Mother, NA24149 [or GM24149]: Father, and NA24385 [or GM24385]: Son), GM11428 (6-year-old female with duplicated chromosome), GM09888 (8-year-old female with trichorhinophalangeal syndrome), GM08331 (4-year-old with chromosome deletion) and GM06226 (6-year-old male with chromosome 1-16 translocation and associated 16p CNV), were obtained from the Bionano Genomics public datasets (https://bionanogenomics.com/library/datasets/). OGM-based genome assembly and variant calling and annotation were performed using Solve version 3.5 (Bionano Genomics). Subsequently, samples were annotated with *nanotatoR* to examine the performance.

Additionally, we tested *nanotatoR*’s ability to accurately annotate the known disease variants in a previously published cohort of 11 Duchenne Muscular Dystrophy samples [27]. For this, internal cohort frequency calculations are based on these 11 samples. For gene expression integration, NA12878 fastq files (RNA-Seq) were obtained from Sequence Read archive (SRA) (Sample GSM754335) and aligned to reference hg19, using STAR [30]. Read counts were estimated using RSEM [31], reported in Transcripts per million (TPM).

### *nanotatoR* input file formats

*nanotatoR* was written in R language. The *nanotatoR* pipeline takes as input Bionano-annotated SV files, in the format of either unmodified SMAP (BNG’s SVcaller output) or text (TXT) files that retain information from the SMAPs, but also append additional fields. The two main differences between the input files are:

#### a) Enzyme

If a combination of restriction endonucleases (Nt.BspQI and Nb.BssSI) are used for DNA labeling, genome assembly and variant calling, the resultant SV call sets from each enzyme are merged into a single TXT file in an SMAP format (SVmerge function, Bionano Genomics). If a single direct labeling enzyme such as DLE1 is used for DNA labeling, the resultant SV call set is kept in a single SMAP file format. Both file types (SVmerge TXT and SMAP) can serve as input files for *nanotatoR*.

#### b) Family

Depending on the availability of family members, the SV-containing input files (TXT/SMAP) contain additional information derived from the *Variant Annotation Pipeline* (BN_VAP, Bionano Genomics). For trio analysis (proband, mother, father), BN_VAP performs molecule checks for SVs identified in the proband (self-molecules) and checks whether the SV is present in the parents’ molecules. For duo analysis (proband vs. control -any family member or unrelated individual- or tumor vs. normal) variants’ presence is evaluated in self-molecules as well as the control sample molecules. For proband-only analysis variants’ presence is evaluated in self-molecules only.

### *nanotatoR* output file formats

The annotations provided by *nanotatoR* are currently subdivided into 5 categories described below: 1) calculation of SV frequency in external and internal databases; 2) determination of gene overlaps; 3) integration of gene expression data; 4) extraction of relevant phenotypic information from public databases for 5) variant filtration. Finally, all of the sub-functions are compiled into a Main function.

### 1. Structural Variant Frequency

Variant frequency is one of the most important filtration characteristics for the identification of rare, possibly pathogenic, variants. Because OGM is not sequence based the average SV breakpoint uncertainty is 3.3 kbp [20]. As a result, compared with SNV frequency calculations, frequency estimates for SVs pose greater difficulty, due to the breakpoint variability between “same” structural variants identified by different techniques.

#### 1.1 External Databases

*nanotatoR* uses 3 external databases: Database of Genomic Variants (DGV) [32], Database of Chromosomal Imbalance and Phenotype in Humans Using Ensembl Resources (DECIPHER) [33] and Bionano Genomics control database (BNDB). The respective functions are named: *DGVfrequency, DECIPHERfrequency* and *BNDBfrequency*. The 3 datasets are accessible through the *nanotatoR* GitHub repository (https://github.com/VilainLab/nanotatoRexternalDB).

BNDB is provided by Bionano Genomics in a subdivided set of 4 files based on the type of SVs (indels, duplications, inversions, and translocations) for two different human reference genomes (GRCh37/hg19 and GRCh38/hg38). *nanotatoR* aggregates the variant files of the user-selected reference genome (hg19 or hg38), into a single format (e.g. TXT) used for frequency calculation. This action is performed as part of the function *BNDBfrequency* with the following input parameters: buildBNInternalDB = TRUE, InternalDBpattern= “hg19” or InternalDBpattern= “hg38”. The following steps are used to calculate the frequency of a query SV in external databases:

##### 1.1a Variant-to-variant similarity

Estimating the frequency of a query SV first requires determining whether the variant is the same as the ones found in a database of interest. In order for the SVs to be considered “same”, *nanotatoR*, by default, checks whether two independent variants of the same type (*e*.*g*. deletion) are on the same chromosome, have 50% or greater size similarity, and if the SV breakpoint start and end positions are within 10 kilobase pairs (kbp) for insertions/deletions/duplications and within 50 kbp for inversions/translocations. For example, if there is a deletion on chromosome 1 with a breakpoint start at position chr1:350,000 and end at chr1:550,000 on the reference, all deletion variants in chr1:340,000-560,000 with a size similarity of 50% would be extracted from the database. Similarly, if the variant was an inversion, *nanotatoR* would search for variants of the same type and on the same chromosome, with a breakpoint start between chr1:300,000 and chr1:400,000 and breakpoint end between chr1:500,000 and chr1:600,000. Currently the 50% size similarity cutoff is not implemented by default for inversions and translocations, as sizes have only started to be provided in the SVcaller output recently; however, users have an option to run the size similarity, and future releases of *nanotatoR* will perform the size similarity calculations by default.

The percentage similarity parameters (DECIPHER and BNDB functions: input parameter perc_similarity; DGV function: input parameter perc_similarity_DGV) and breakpoint start and end error (DECIPHER and BNG functions insertion, deletion and duplication: input parameter “win_indel”; DGV function insertion, deletion and duplication: win_indel_DGV; DECIPHER and BNG functions inversion and translocation: win_inv_trans; DGV function inversion and translocation: win_inv_trans_DGV) are modifiable by the user.

##### 1.1b Variant size and confidence score

Two additional criteria are implemented to select for high-quality variants in BNDB. Bionano’s SVcaller calculates a confidence score for insertions, deletions, inversions, and translocations. To calculate allele frequency, *nanotatoR* takes into account the BNDB variants above a threshold quality score of 0.5 for insertions and deletions (indelconf), 0.01 for inversions (invconf) and 0.1 for translocations (transconf). These thresholds can be modified by the user. In addition, *nanotatoR* filters out SVs below 1 kbp in size to decrease the likelihood of false positive calls [20].

##### 1.1c Zygosity

Variants in BNDB are reported as homozygous, heterozygous or “unknown” (DGV and DECIPHER do not report zygosity). This is used to refine frequency calculation for BNDB SVs: *nanotatoR* attributes an allele count of 2 for homozygous SVs and 1 heterozygous SVs. Currently, *nanotatoR* overestimates the frequency for variants that overlap with reference database (BNDB) SVs for which the zygosity is unknown by counting the number of alleles as 2. If the query SV matches with multiple variants in the BNDB from the same BNDB sample, *nanotatoR* counts these as a single variant/sample, with allele count of 2 for homozygous/unknown and 1 for heterozygous matches.

##### 1.1d Frequency calculations

For DECIPHER and DGV, SV frequency is calculated by dividing the number of query matched database variants (step 1.1a) by the total number of alleles in the database, *i*.*e*. 2x the number of samples, which are diploid, and multiplying with 100 to get percentage frequency (Formula 1).

**Formula 1: External public (DGV and DECIPHER) database SV frequency calculation**. Numerator: number of variants that pass the similarity criteria (step 1.1.a). Denominator is twice the number of samples, *i*.*e*. the number of alleles. This ratio is multiplied by 100 to express the frequency as a percentage.

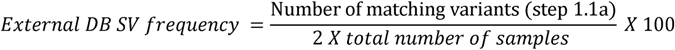

For BNDB two types of frequency calculations are performed: filtered and unfiltered. For filtered frequency calculations the following criteria must be met: 1.1a; 1.1b; 1.1c. For unfiltered variants frequency calculation only 1.1a and 1.1c criteria are enforced. The resultant number of identified counts is divided by the number of alleles in BNDB (currently 468 for 234 diploid samples). The result is multiplied by 100 to get a percentage (Formula 2).

**Formula 2: BNDB database filtered SV frequency calculation**. The variants that pass the similarity criterion (step 1.1.a) are filtered with size threshold and quality score (step 1.1.b). The number of variants is estimated as mentioned in step 1.1.c. Denominator is the number of alleles. This ratio is multiplied by 100 to get the frequency in percentage.

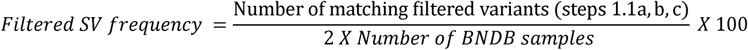

###### Output

The output is appended to the original input file in individual columns. For DECIPHER, this consists of a single column termed “DECIPHER_Freq_Perc”. As DGV provides information on number of samples in addition to frequency, *nanotatoR* prints two columns: “DGV_Count” (with the total number of unique DGV samples containing variants matching the query SV) and “DGV_Freq_Perc” (for the percentage calculated using Formula 1). For the BNDB, in addition to “BNG_Freq_Perc_Filtered”, “BNG_Freq_Perc_UnFiltered”, a third column reports “BNG_Homozygotes” (number of homozygous variants that pass the filtration criteria).

#### 1.2 Internal Databases

The internal cohort analysis is designed to calculate variant frequency based on aggregation of SVs for samples ran within an institution or laboratory and provides parental zygosity information for inherited variants in familial cases. The function consists of two distinct parts:

##### 1.2a Building the internal cohort database

Individual (solo) SMAP files for each of the samples are concatenated to build an internal database (buildSVInternalDB = TRUE), which is stored in the form of a text file. This step creates a unique sample identifier (nanoID) based on a key provided that ensures unique sample ID and encodes family relatedness. The nanoID is written as *NR<Family #>*.*<Relationship #>*. For example, the proband in a family of three (trio) would be denoted as NR23.1, with *NR23* denoting the family ID and *1* denoting the proband. For the parents of this proband, the nanoID would be NR23.2 for the mother and NR23.3 for the father. Currently, only trio analyses are supported, future updates will include larger family analyses. If multiple projects exist within the same institution and are coded with project-specific identifiers *nanotatoR* will append the project-specific identifier in front of the nanoID (*e*.*g*. *Project1*_NR23.1 and *Project2*_NR42.1).

##### 1.2b Calculating internal frequency and determining parental zygosity

For singleton analyses, the function *internalFrequency_Solo* (for both DLE labeling and SVmerge) calculates internal database frequency of queried SVs based on the same principles explained in section 1.1d (Formula 2) for BNDB frequency calculations. However, additional filtration criteria are implemented to increase the accuracy of frequency estimation. SVs overlapping gaps in hg19/hg38 are annotated in the output SMAPs as “nbase” calls (*e*.*g*. “deletion_nbase”) and are likely to be false. *nanotatoR* filters out “nbase”-containing SVs when estimating internal frequency. For duplications, inversions, and translocations *nanotatoR* evaluates whether chimeric scores “pass” the thresholds set by the Bionano SVcaller during *de-novo* genome assembly [34] ensuring that SVs that “fail” this criterion are eliminated from internal frequency calculations (Fail_BSPQI_assembly_chimeric_score = “pass” or Fail_BSSSI_assembly_chimeric_score = “pass”) for SVmerge datasets, or (Fail_assembly_chimeric_score = “pass”) for a single-enzyme dataset. Lastly, *nanotatoR* checks whether the SVs were confirmed with Bionano Variant Annotation Pipeline, which examines individual molecules for support of the identified SV [34] (Found_in_self_BSPQI_molecules = “yes” or Found_in_self_BSSSI_molecule = “yes”) for SVmerge datasets, or (Found_in_self_molecules = “yes”) for a single-enzyme dataset.

For family analyses (duos and trios), the *internalFrequencyTrio_Duo* function is used to identify parental/control sample zygosity based on the nanoID coding using criteria described in sections 1.1.a/c (note that, here, the default size similarity percentage used is ≥ 90% as inherited variants are expected to be virtually identical). Zygosity information for the identified variants is extracted and appended into two separate columns (fatherZygosity and motherZygosity). This functionality is available for both SVmerge (merged outputs from 2 enzymes) and single enzyme labeling. SVs with the same family ID as the query are not included in the overall internal frequency calculation as described in the previous paragraph. Five columns: “MotherZygosity”, “FatherZygosity”, “Internal_Freq_Perc_Filtered”, “Internal_Freq_Perc_Unfiltered”, and “Internal_Homozygotes” are appended to each of the annotated input files (nonrelevant fields contain dashes).

### 2. Gene Overlap

The gene overlap function (*overlapnearestgeneSearch*) identifies known gene and non-coding RNA genomic locations that overlap with or are immediately upstream or downstream from the identified SVs. This function takes as input an SV-containing file (TXT or SMAP) and a modified Browser Extensible Data (BED) file where human X and Y chromosomes are numbered as 23 and 24 respectively. The user has an option of either providing a Bionano-provided modified BED file (inputfmtBed = “BNBED”) or a BED file from UCSC Genome Browser or GENCODE (inputfmtBed = “BED”). For the latter, *nanotatoR* supports conversion of a BED file into BNBED standard with *buildrunBNBedFiles* function. The BED file is used to extract location/orientation of genes and overlap this information with SVs (*overlapGenes* function, called from *overlapnearestgeneSearch*). The function calculates the percentage overlap, by calculating its distance from the breakpoint start (if the gene is partially upstream of the SV) or breakpoint end (if the gene is partially downstream of the SV), and dividing it by the length of the gene (calculated by *nanotatoR* from genomic coordinates information in the BED file). By default, *nanotatoR* applies a 3 kbp gene overlap window (breakpoint start – 3 kbp; breakpoint end + 3 kbp) to account for the typical OGM breakpoint error [20] when searching for genes overlapping insertions, deletions and duplications. For inversions and translocations overlapping genes are limited to +/−10 kbp from the breakpoint start/end (both parameters are user-selectable). For the *nonOverlapGenes* function (also called from *overlapnearestgeneSearch)*, genes located near, but not overlapping with, SVs are reported along with the corresponding distances from the SV. Genes are sorted based on their distance from the SV breakpoints. The default number of reported nonoverlap genes is 3 (also user-selectable). The output produces 3 additional columns: “OverlapGenes_strand_perc”, “Upstream_nonOverlapGenes_dist_kb” and “Downstream_nonOverlapGenes_dist_kb”.

### 3. Expression Data Integration

The *SVexpression_solo/_duo/_trio* functions for singletons, dyads and trios respectively provide the user with tools to integrate tissue-specific gene expression values with SVs. The function takes as input a matrix of gene names and corresponding expression values for each sample (individual files can be merged by the *RNAseqcombine* function for dyads/trios or *RNAseqcombine_solo* function for singletons). To differentiate between sample types (probands and affected/unaffected parents), we recommend the user add a code (or pattern) to the file name. For example, for the proband of family 23, the expression file name would be Sample23_P_expression.txt, where P denotes proband. Currently, by default *nanotatoR* recognizes “P” as proband, “UM/AM” for unaffected/affected mother and “UF/AF” for unaffected/affected father. A function to encode more complex intra-family relatedness and identify individual samples, termed nanoID, is in development and will be available by default in the next release of *nanotatoR*.

Expression values for overlapping and non-overlapping SV genes are extracted from the genome-wide expression matrix and appended into separate columns in the overall SV input file. For example, an overlap of gene X with a SV in the proband, with an expression value of 10, would be represented as gene X (10), and printed in the “OverlapProbandEXP**”** column. The appended number of columns in the output is dependent on which functions were run. For a trio analysis, 9 columns are added: “OverlapProbandEXP”, “OverlapFatherEXP”, and “OverlapMotherEXP” for overlapping genes, and a similar set for up- and down-stream non-overlapping genes (*e*.*g*. “NonOverlapUPprobandEXP”, “NonOverlapDNprobandEXP”). In case of dyads the number of columns would be 6 (with the parent column being either mother or father) and 3 for singletons.

### 4. Entrez Extract

The *gene_list_generation* function assembles a list of genes based on the patient’s phenotype and overlaps it with gene names that span SVs. User-provided, phenotype-based keywords are used to generate a gene list from the following databases: ClinVar [35], OMIM (https://omim.org/), GTR [36], and the NCBI’s Gene database (www.ncbi.nlm.nih.gov/gene). The input to the function is a term, which can be provided as a single term input (method = “Single”), a vector of terms (method = “Multiple”), or a text file (method = “Text”). The output can be a dataframe or text. The *rentrez* [37] and *VarfromPDB* [38] R-language packages are used to extract data related to each of the user-provided phenotypic terms, from the individual databases. For the Gene database *rentrez* provides the entrez IDs associated with each gene, which are converted in *nanotatoR* to gene symbols using org.Hs.eg.db [37], a Bioconductor package. For OMIM, *rentrez* provides the OMIM record IDs, which are used to extract the corresponding disease-associated genes from the OMIM ID-to-gene ID conversion dataset (mim2gene.txt). For GTR, *rentrez* extracts the GTR record IDs, which are then used to extract corresponding gene symbols from the downloaded GTR database. For ClinVar, *VarfromPDB* is used to extract genes corresponding to the input term. All genes to which the query keyword is attached, irrespective of their clinical significance, are extracted; genes of clinical significance (*i*.*e*. those for which Pathogenic/Likely Pathogenic variants are reported) are further reported in a separate column. The user also has the option to download the ClinVar and GTR databases by choosing downloadClinvar = TRUE and downloadGTR = TRUE, which may improve run times. The user has an option to save the datasets (removeClinvar = FALSE and removeGTR = FALSE) or delete the database after the analysis is completed (removeClinvar = TRUE and removeGTR = TRUE).

The output is provided in CSV format with 3 columns: “Genes”, “Terms”, and “ClinicalSignificance”. The “Terms” column contains the list of terms associated to each gene and corresponding database from where the association was derived. The “**ClinicalSignificance**” column contains genes that have clinical significance (Pathogenic/Likely Pathogenic variants) for the associated term, derived from the ClinVar database. The output of *entrez* extract serves as input for the subsequent variant filtration step.

### 5. Variant Filtration

The filtration function has two major functionalities: categorization of variants into groups (such as *de novo*, inherited from mother/father; potential compound heterozygous; inversions or translocations) and integration of the primary gene list (either provided by the user or generated by *nanotatoR* as described in section 4) into the input SV-containing file. In this function, genes overlapping and near SVs that are present in the primary gene list are printed in separate columns. For genes that overlap with a SV, the output is printed in columns termed “Overlap_PG**”** (primary genes that are common with the genes that overlap SVs) and “Overlap_PG_Terms**”** (terms and database the genes are being extracted from). Genes that are up/downstream of, but not overlapping with, the SV are divided into “Non_Overlap_UP_PG”, “Non_Overlap_UP_Terms”, “Non_Overlap_DN_PG”, and “Non_Overlap_DN_Terms”.

The final annotated SV calls are divided into multiple sheets:

- *All*: contains all identified variant types and annotations.
- *all_PG_OV*: Variants overlapping with the primary gene list.
- *Mismatch*: Variant Type = “MisMatch” in SVmerge output (available for dual enzyme labeling output only) contains SV calls discordant between the two enzymes datasets (*e*.*g*. one called a deletion and the second an insertion).

By default, for the rest of the sheets, *nanotatoR* performs self-molecule checks for all SV types (described in section 1.2b, Found_in_self_ molecules = yes) and chimeric score quality checks for duplications, inversions and translocations (also described in section 1.2b, Fail_BSPQI_assembly_chimeric_score **= “**pass”) and reports on zygosity. (These two criteria are henceforth called “default *nanotatoR* filtration”). For indel_dup, this comprises:

- For singleton studies (solo): Variant type = “insertion”, “deletion”, “duplication”, “duplication_split” or “duplication_inverted”.
- For dyads: indel_dup_notShared or indel_dup_Shared respectively: Insertions, deletions and duplications not present in a control sample, mother or father (Found_in_control_molecules **= “**no”) or present in a control sample, mother or father (Found_in_control_molecules **= “**yes”).
- For trios, insertions, deletions or duplications are reported in:

⋄indel_dup_denovo if not present in the parents (Found_in_parents_BSPQI_molecules/Found_in_parents_BSSSI_molecules **=** “none” or “-” for SVmerge or Found_in_parents_molecules **=** “none” for single enzyme).
⋄indel_dup_both if present in the proband, as well as both parents (*e*.*g*. Found_in_parents_molecules **=** “both”).
⋄indel_dup_mother and indel_dup_father if inherited from only the mother or father, respectively (*e*.*g*. Found_in_parents_molecules **=** “mother” or “father”).
⋄indel_dup_cmpdHET if present in the heterozygous state in parents. This output is a combination of Indel_dup_mother and Indel_dup_father datasets. The user can manually inspect this list to find potential compound heterozygote variants, *i*.*e*. 2 variants with genomic coordinates overlapping with the same gene, one present in the mother, the other in the father.

Finally, inversion and translocations are reported as follows, for singletons, dyads, or trios:

- Variant Type = “inversion” or “inversion_paired” or “inversion_partial” or “inversion_repeat”.
- Variant Type = “translocation_intrachr” or “translocation_interchr” if within a single chromosome or between two chromosomes respectively.

There is a total of 6 variant filtration functions, based on enzyme type (SVmerge for dual-enzyme labeling or single enzyme) and sample type (singleton, dyad, or trio). For example, for a single-enzyme singleton dataset the function is *run_bionano_filter_SE_solo*. Others are *run_bionano_filter_SE_duo, run_bionano_filter_SE_trio, run_bionano_filter_SVmerge_solo, run_bionano_filter_SVmerge_duo*, and *run_bionano_filter_SVmerge_trio*. Variant filtration output is an Excel file with each of the different output groups represented as separate tabs. The output is written in Excel file format using the openxlsx [39] package, with the following default naming convention “Sample1_solo.xlsx”.

### Main Function

The main function sequentially runs the available *nanotatoR* sub-functions by merging the outputs from each step. There are in total 6 main functions depending on the type of input: *nanotatoR_main_Solo_SE; nanotatoR_main_Duo_SE; nanotatoR_main_Trio_SE; nanotatoR_main_Solo_SVmerge; nanotatoR_Duo_SVmerge and nanotatoR_SVmerge_Trio*. The Main function takes the SMAP file, DGV file, BED file, internal database file and phenotype term list as inputs and provides a single output file in a form of an Excel spreadsheet. The output location and file name are user-specified.

## Results

The *nanotatoR* pipeline, written in R, integrates five individual sub-functions based on the enzyme and the sample type. A visual representation of the steps is shown in **Figure 1**. *nanotatoR* passed all runtime and space criteria for the Bioconductor repository (https://www.bioconductor.org/developers/package-guidelines/), where it was accepted in the April, 2019 cycle. The Bioconductor link for *nanotatoR* is https://bioconductor.org/packages/devel/bioc/html/nanotatoR.html, and the latest version update is available at *https://github.com/VilainLab/nanotatoR*.

**Figure 1.**
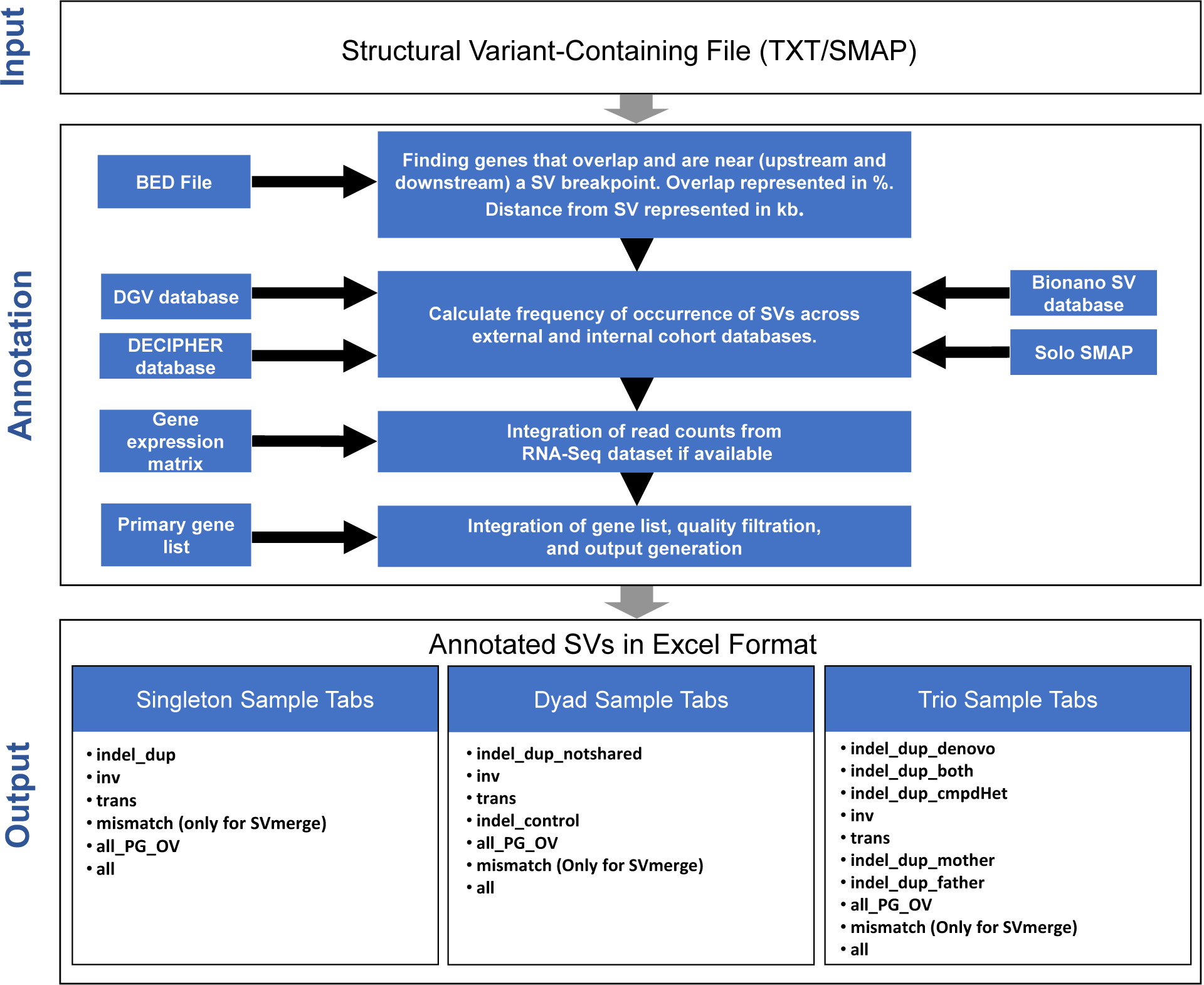
Workflow of the *nanotatoR* pipeline: The nanotatoR pipeline is divided into 3 layers. **Input Layer:** Takes as input the OGM text or Smap file. **Annotation Layer**: The annotation layer comprises of five methods. The method that extracts the overlapping genes and the genes near (downstream and upstream) takes as input a BED file, and calculates the overlap percentage and the distance between nearest genes and SV using chromosomal locations. Next, the frequency calculation function calculates external and internal frequency taking DGV, DECIPHER and BNDB database as input for external frequency calculation, while input solo files, merged to form the internal frequency database, are taken as input to calculate the internal frequency. If RNA-Seq data is available, the expression count matrix is taken as input. Finally, output from all these methods as well as a primary gene list created from terms, is integrated, filtered based on quality criteria and written into an Excel file. **Output Layer:** The output is an Excel workbook, with each tab representing different SV types. The output files and number of tabs depend on the sample type and enzyme type: Singleton samples have 5 tabs for DLE and 6 tabs for SVmerge; 6 tabs for DLE and 7 tabs for SVmerge are created for dyad analyses; Trio analyses have 9 tabs for DLE and 10 tabs for SVmerge.

The output is in the form of an Excel workbook subdivided into variant types and inheritance modes in familial cases. The user has an option to either filter the data based on input parameters or perform the filtrations steps in the final Excel sheets. Theoretical examples of the *nanotatoR* annotation process and output are illustrated in **Figure 2** for various types of SVs.

**Figure 2:**
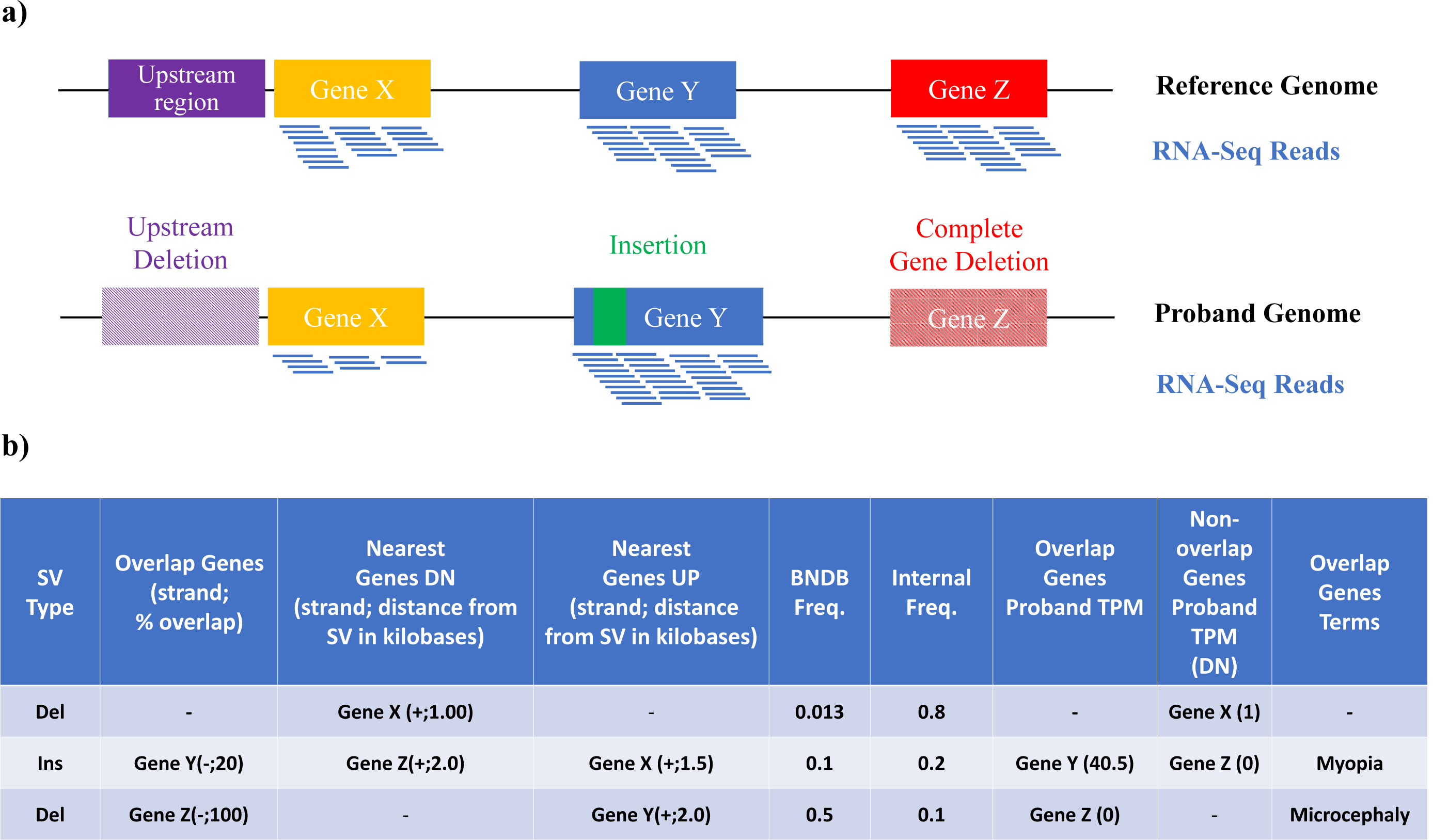
*nanotatoR* annotates genes overlapping or near a SV: **a) The cartoon shows three hypothetical scenarios:** one deletion in the region upstream of Gene X (yellow) which may contain regulatory regions, indicated as solid purple in the reference genome (top) and striped in the patient’s genome (bottom); one insertion (green) into Gene Y (blue), and a complete deletion of Gene Z (red, striped in patient genome). RNA-Seq reads are depicted as blue lines below the genes. **b) *nanotatoR* annotation snapshot:** *nanotatoR* annotates the three variants with the overlapping genes and percentage overlap, nearest genes upstream and downstream, distance to the breakpoints in kilobases, BNDB frequency, internal frequency, overlap gene expression value (in transcripts per million or TPM), nearest genes expression in TPM and overlapping genes term from NCBI databases.

To demonstrate the various functionalities of *nanotatoR*, we present below the annotation results obtained from previously described truth sets: a control trio mapped with the single-enzyme technique, a control singleton sample mapped with both DLE and two-enzyme techniques, and a cohort of patients, for which we have previously established the efficacy of OGM to identify the SV causing Duchenne Muscular Dystrophy [27].

### Example I: Annotation of a control trio single labeling dataset

We used the so-called “Ashkenazi trio” reference datasets, mapped using the DLE labeling methodology, to test the trio analysis function of *nanotatoR* (expression data was not available for these samples). Bionano SVcaller identified a total of 9,387 SVs in the proband (NA24385), shown in the last tab of the 9-tab report, termed “*all*”. The complete, unfiltered *nanotatoR* output file for NA24385 (GM24385) is available in **Supplementary Table S1**.

#### nanotatoR filtration and variant type annotation

After *nanotatoR* filtration (method described in section 5 Variant Filtration of Materials and Methods section), 8,804 variants (93.8%) remained of the original 9,387. The variants were annotated based on criteria described in Methods section 5. SVs were distributed as shown in **Figure 3** (left pie chart). The vast majority (8,680) were in the *“indel_dup”* tabs; 114 (out of 279 unfiltered) inversions were reported in the “*inv*” tab, and 0 (out of 84) translocations in the “*trans*” tab. All the translocations called by the Bionano SVcaller in this sample were in the categories “trans_interchr_common” and “trans_intrachr_common”, which are classified as likely false by the Bionano annotation pipeline [34, 40].

**Figure 3.**
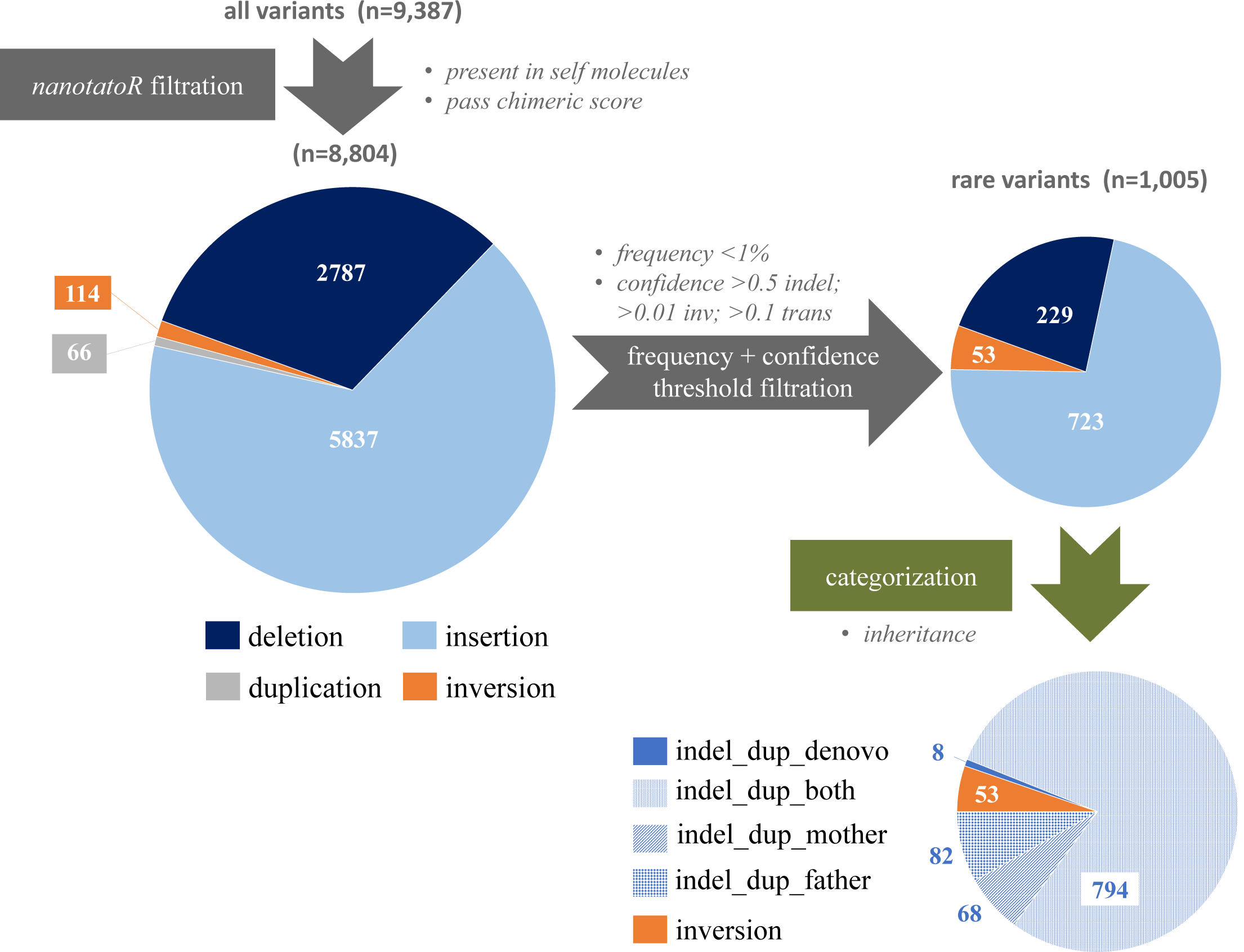
Filtration and annotation of SV distribution in the NA24385 trio dataset: Out of the total 9387 variants found by SVcaller, 8,804 passed the *nanotatoR* filtration of “Present in self molecules” and “Pass chimeric score” conditions. Of these, 2,787 were deletions (dark blue), 5,837 were insertions (light blue), 66 were duplications (grey), 114 were inversions (orange) and 0 were translocations (not shown), left pie chart. Note, *nanotatoR* outputs deletions, insertions and duplications in a single excel sheet “indel_dup”. The number of variants was dramatically reduced after filtering for rare variants and imposing a confidence threshold (top right pie chart). With a threshold of less than 1% internal frequency, DGV frequency, BNDB frequency, and DECIPHER frequency and confidence thresholds of >0.5 for INDELs, >0.01 for inversions and >0.1 for translocations, 1,005 rare variants remain. These were further categorized with *nanotatoR* by inheritance (bottom right pie chart). All 53 inversions were inherited. Of the 952 indel_dup variants, only 8 were *de novo*, 794 are were identified as indel_dup_both (found in both mother and father), 68 are indel_dup_mother (found in only the mother), and 82 are indel_dup_father. This annotation can be used to evaluate relevance of the variants to the condition studied.

#### Confidence and frequency filtration

To demonstrate the importance of frequency filtration in identifying rare variants, a manual filtration was performed using a 1% threshold for external databases (DGV, DECIPHER and BNDB) and the internal cohort database. A confidence threshold (0.5 for insertions and deletions, 0.01 for inversions) was also applied. This eliminated 89% of the variants, leaving a total of 1,005 SVs, including 53 inv variants and 952 indel_dup (shown in top right pie chart, **Fig. 3**).

#### Inheritance annotation

(**Fig. 3**, bottom right pie chart): All inversion variants were inherited. Out of the 952 insertions, deletions and duplications, only 8 were *de novo* (“*indel_dup_denovo*” tab), 794 found in both parents (“*indel_dup_both*”), 68 in the mother only (“*indel_dup_mother*”), and 82 in the father only (“*indel_dup_father*”). These numbers can be used to evaluate pathogenicity of the SVs. 150 SVs would be reported in the “*indel_dup_cmdHet*” column (found in either the mother or father, but not both), which can be manually inspected to identify potential compound heterozygous SVs.

#### SVs overlapping with the primary gene list

As these samples are those of healthy individuals, we could not use a disease term to generate a gene list. However, analysis of the genomes with OGM had revealed a deletion variant affecting the *UGT2B17* gene in the son and the mother [41]. A 150 kb deletion on chromosome 4q13.2 spanning the whole *UGT2B17* gene has been associated with osteoporosis [42]. To check whether our tool can efficiently annotate the variant, we used the term ‘osteoporosis’ to generate a primary gene list. This yielded a list of over 307 genes of which only 4 had pathogenic or likely pathogenic variants in ClinVar, highlighting the importance of this *nanotatoR* function (**Supplementary Table S2**). The complete extracted gene list can be used for gene discovery, while the pathogenic list is most efficient to identify variants in genes known to be associated with the proband’s phenotype. Note that while *UGT2B17* was accurately extracted into the primary gene list by *nanotatoR*, as its association with osteoporosis is reported in OMIM, it does not appear in the list of genes with pathogenic variants, as no such variant is currently reported in ClinVar.

169 SVs were found to be overlapping with the primary gene list, and were shown in the “*all_PG_OV*” tab.

A deletion in *UGT2B17* gene is observed in both the “*indel_dup_mother*” tab and the “*all_PG_OV*” tab, as expected [41]. The SV overlaps with the *UGT2B17* gene and 4 pseudogenes UGT2B29P, AC147055.2, AC147055.3 and AC147055.4 as illustrated in **Figure S1**.

#### Internal cohort frequency and zygosity calculations for the UGT2B17 variant

To investigate the frequency of the variant in the 8-sample internal cohort database, we first selected all variants with within −10 kb of start breakpoint and +10 kb of the end breakpoint (*i*.*e*. between hg19 genomic coordinates chr4:69,362,091 and chr4:69,500,860). A total of 6 variants passed the filtration criteria (“*GM24385_del_totalData*” tab in **Table S3**). Of these, 3 are from the query family: one is the proband’s variant (all annotation shown in “*GM24385_Variant_UGT2B17*” tab in **Table S3**) and the other two are found in his mother. Of the maternal variants, one has exactly same start and end breakpoints as the proband’s. Bionano SVcaller also called another variant in the mother with the same end breakpoint, and a similar, but not identical, start breakpoint. Both were retained as they have a size similarity > 90% with the proband’s variant. As described in Methods section 1.2b, *nanotatoR* selects the variants that pass size similarity and breakpoint criteria, and reports the zygosity for each in the parents (“*GM24385_del_example_Zygosity”* tab in **Table S3**).

In addition to the family, the variant was found in two other samples of the internal cohort. The first was heterozygous, the second homozygous, so the total number of alleles carrying the SV was counted as 3. As the internal control cohort was composed of 8 samples, of which 3 were part of the Ashkenazi family, the total number of alleles in the internal cohort was calculated as 10 = 2 × (8-3), where 2 is for diploid genomes, 8 is the total number of samples in the cohort and 3 is the number of related individuals. The final internal frequency thus is (3/10) *100 = 30% (for both filtered and unfiltered, as all variants passed the quality filters). Note that the *nanotatoR* annotation process has detected the erroneous duplicate call made by SVcaller in sample NA12878, where two variants with identical characteristics were called under two different SVIndex numbers (rows 6&7, totalData tab, Table S3). Only one is taken into account for internal frequency calculation, as shown in **Table S3**, where tabs “*GM24385_del_example_filter”* and “*GM24385_del_example_unfilt*” show the samples used for the calculations.

Next, we calculated the filtered and unfiltered frequency (Formula 2, Methods 1.1d) of the deletion overlapping *UGT2B17* in the Bionano reference database. We identified a total of 58 variants in BNDB (“*GM24385_data_all”* tab in **Supplementary Table S4**). Of these 33 (“*GM24385_data_filtered”* and “*GM24385_data_unfiltered”* tabs of **Supplementary Table S4)** passed the *nanotatoR* default filtration criteria and were used for frequency calculation. Twelve were homozygotes and 21 heterozygotes for a total number of variant alleles of 43. The total number of samples in BNDB is 234, hence the variant frequency is calculated as (43/468) *100 = 9.18%. (Note that, here too, the number of variants was the same before and after filtration, yielding the same frequency value).

#### Run times

The SMAPs were annotated on an Intel core i7-6700 CPU with 16 GB RAM, Windows 10 system. It took ∼15 minutes to annotate the trio sample (**Supplementary Table S5** has run time for each of the functions individually). The runtime for *nanotatoR* is dependent on number of variants as well as network speed (for *gene_list_generation* function).

To generate the primary gene list for the trio sample, we downloaded the ClinVar and GTR databases, using the downloadClinvar = TRUE and downloadGTR = TRUE parameters in the *gene_list_generation* function. The *gene_list_generation* function took ∼2 minutes to run for the sample. The time for this function is dependent on the number of input terms, as well as the computational/internet bandwidth available to the user. To make this process faster, the input parameters *removeGTR* and *removeClinvar* can be switched to FALSE; *nanotatoR* will then use the pre-downloaded database files for subsequent runs. It is recommended to download these databases periodically as they get updated frequently.

All the other databases (OMIM, DGV, DECIPHER, BNDB) must be downloaded manually from the database websites or from the *nanotatoR* database GitHub page (https://github.com/VilainLab/nanotatoRexternalDB). For this example, the internal frequency database was built based on a cohort of 8 samples (time ∼10 minutes). The time taken for database creation depends on the user’s cohort size.

### Example II: Annotation of a control singleton dataset labeled with several enzymes

We also investigated the OGM datasets available for the sample NA12878 to test the annotation effectiveness of *nanotatoR* on multi-enzyme labeling and integration of RNA-Seq data. OGM data is available for three labeling enzymes Nt.BspQI, Nb.BssSI and DLE1. The Nt.BspQI and Nb.BssSI SMAP outputs were merged using SVmerge. 10,087 and 6,814 SVs were reported by SVcaller for single enzyme (DLE1) and SVmerge output respectively (**Figure 4**).

SV annotation for a single enzyme took approximately 28 minutes and 24 minutes for SVmerge data output. The time taken for each of the functions is reported in **Supplementary Table S5**. Currently, the expression data aggregation function takes the longest time as it extracts each of the genes overlapping the SVs and finds the corresponding expression values from the RNA-Seq datasets. The run time for this function largely depends on the number of called SVs and the number of genes that overlap with them. The complete *nanotatoR*-annotated output Excel files can be found in **Supplementary Tables S6 and S7** for DLE1 labeling and SVmerge (Nt.BspQI/Nb.BssSI) respectively.

**Figure 4.**
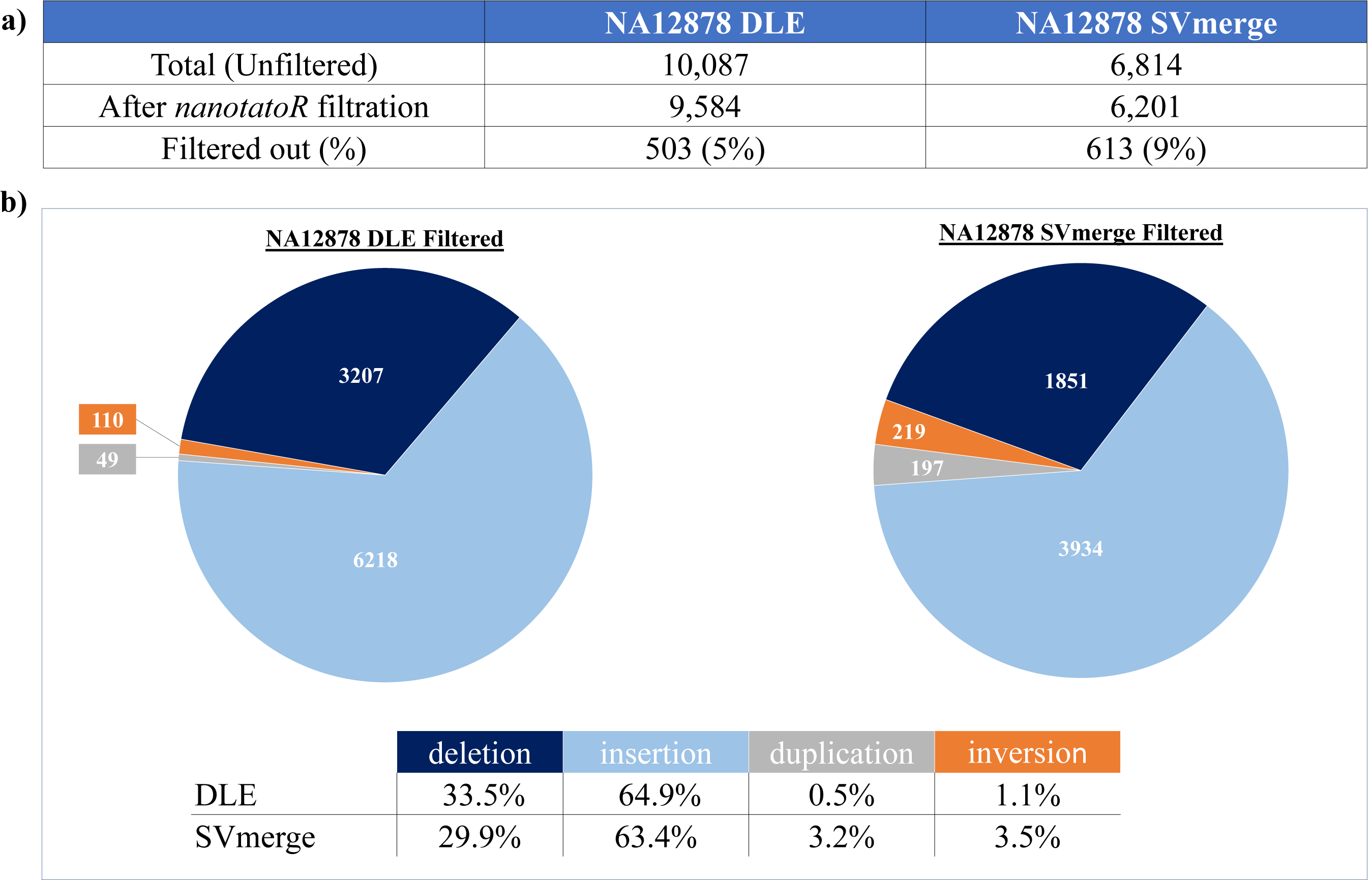
Variant distribution for the singleton NA12878 sample. **a) Unfiltered and filtered variants distribution in DLE and SVmerge datasets:** The total number of unfiltered variants for NA12878 DLE are 10,087, out of which 9,584 variants are filtered using *nanotatoR* criteria (found in self molecules, passed chimeric score threshold). For NA12878 SVmerge, out of 6,814 variants, 6,201 pass the filtration. Mismatches were not considered in this analysis but are shown in Table S7. **b) SV distribution in the NA12878 DLE and SVmerge filtered datasets:** Deletions (dark blue), insertions (light blue), duplications (grey) and inversions (orange) numbers are as shown in the pie charts. Bottom table shows distribution of SVs by type in percentages. For DLE, the majority of the identified SVs were insertions (64.9%), followed by deletions (33.5%), inversions (1.1%) and finally duplications (0.5%). While the total number of variants called is different between DLE and SVmerge, a similar pattern is seen in the SVmerge dataset. Many more duplications and inversions were called in the dual labeling than single DLE labeling method.

For DLE1 labeling, out of the 10,087 variants called by the SVcaller, 9,584 (∼5%) remained after default *nanotatoR* filtration (“Present in self molecules” and “Pass chimeric score”, see Methods section 5) (**Figure 4a**). As for the trio genomes, the vast majority of called SVs were indels, with a similar number of inversions (110) and zero translocations. Further breakdown of the *indel_dup* tab reveals 3,207 deletions, 6,218 insertions, and 49 duplications (**Figure 4b**; **left panel**). For dual-enzyme labeling, fewer variants were called in the SVmerge dataset (6,814), of which ∼9% were filtered out by *nanotatoR* default filtration. Of the remaining 6,201, 5,982 are “*indel_dup”*, 219 “*inv”*, 0 “*trans”* and 11 *mismatches*. Breakdown of the indel SVs between deletions, insertions and duplications is shown in **Fig. 4b, right panel)**. Proportions of insertions and deletions are similar in the two data sets, while the dual enzyme labeling called more duplications and inversions than single-enzyme labeling in this example.

To validate the efficiency of *nanotatoR* in identifying genes overlapping with SVs for NA12878, we looked for four previously published variants [43]. The 4 deletion variants identified in the study overlapped *GSTM1, LCE3B, LCE3C, CR1* and *SIGLEC14* genes. *nanotatoR’*s automated pipeline was able to identify the same type of variant (deletions) involving the same genes in both the single enzyme and SVmerge datasets. SV breakpoints reported in the original publication and in the *nanotatoR*-annotated data sets are shown in **Supplementary Table S8**; SV type and gene names are highlighted in the *all_PG_OV* tab in **Tables S6** (DLE) and **S7** (SVmerge).

### Example III: Duchenne Muscular Dystrophy Cohort

We have previously published validation of the OGM technology to identify variants in the DMD gene in a cohort of patients with Duchenne muscular dystrophy [27]. We used the same cohort to test the *nanotatoR* annotation pipeline. The *gene_list_generation* function, using “Duchenne muscular dystrophy” as input for the *rentrez* tool, used at high stringency, *i*.*e*. selecting only genes with pathogenic or likely pathogenic variants in ClinVar, yielded only one gene as expected for this monogenic disorder. **Table 1** shows that all of the previously identified variants in DMD cases were correctly annotated. Each of these types of variants was placed in the correct final Excel output tab with corresponding frequencies, gene overlap and maternal carrier status. (Annotation for all samples is shown in **Supplementary Table S9**; columns used to create Table 1 are highlighted).

**Table 1:** Summary of *nanotatoR* annotation results of Duchenne muscular dystrophy patient cohort

| Sample ID | Overlap Gene | Variant Type | Clinical Significance | Zygosity | Internal Frequency | DGV/BNDB Frequency |
| --- | --- | --- | --- | --- | --- | --- |
| CDMD1003_P | *DMD* | Deletion | Pathogenic | Hemizygous | 0 | 0/0 |
| CDMD1155_P | *DMD* | Deletion | Pathogenic | Hemizygous | 0 | 0/0 |
| CDMD1156_P | *DMD* | Deletion | Pathogenic | Hemizygous | 0 | 0/0 |
| CDMD1159_P | *DMD* | Deletion | Pathogenic | Hemizygous | 0 | 0/0 |
|  | *DMD* | Deletion | Unknown | Hemizygous | 0 | 0/0 |
| CDMD1131_P<br>CDMD1132_M | *DMD* | Deletion | Pathogenic<br>Carrier | Hemizygous<br>Heterozygous | 22% | 0/0 |
| CDMD1157_P<br>CDMD1158_M | *DMD* | Deletion | Pathogenic<br>Non-Carrier | Hemizygous<br>n/a | 11% | 0/0 |
| CDMD1163_P<br>CDMD1164_M | *DMD* | Insertion | Pathogenic<br>Carrier | Hemizygous<br>Heterozygous | 0 | 0/0 |
| CDMD1187_P | *DMD* | Inversion | Pathogenic | Hemizygous | 0 | 0/0 |

Using *nanotatoR* we annotated variants that overlapped *DMD* gene, evaluated the zygosity and calculated the internal (cohort size 11 samples) and external (DGV/BNDB) frequencies. The details of the variants can be found in Barseghyan et.al. 2018.

Note that, as most callers, Bionano SVcaller currently identifies zygosity for X and Y chromosome variants in an XY individual as homozygous rather than hemizygous, which we corrected in Table 1. As a result of the erroneous input, internal frequency is currently overestimated for variants on the X chromosome. Frequency calculations for SVs on the Y chromosome are not affected.

Using *nanotatoR*, we were able to automate steps that previously had to be taken manually to identify the pathogenic SVs in the *DMD* gene in the data sets: navigation to X chromosome location of the *DMD* gene, selection of the type of the SV (deletion/insertion, etc..), filtration by frequency, and curation of gene pathogenicity. In addition to the previously reported SVs, with the help of *nanotatoR*, we identified an additional deletion of unknown significance in sample CDMD_1159 that had previously been missed.

## Conclusions

Structural variants play a major role in various genetic diseases (reviewed in [44]). Due to the technical limitations of short-read-based genome sequencing and microarray techniques, identification of SVs is challenging. Introduction of optical genome mapping and long-read-based technologies promises to advance the field of SV identification. Although there are tools available for annotation of SVs (AnnotSV [45], Annovar [46]), they do not take into account OGM criteria (like self molecules or chimeric score) for filtration.. This has prompted the development of *nanotatoR* to help researchers analyze OGM SV datasets, with high efficiency and precision. The annotation pipelines available for the OGM and LRS data are currently suboptimal, with limited user-defined parameters for frequency calculations in external databases, intersection with gene expression datasets, or filtration through primary gene lists. These functions are critical for clinical applications to evaluate SV pathogenicity. *nanotatoR*, currently takes as input the SMAP/TXT variant file, databases (internal and/or external), terms list, gene location bed files, and expression values, to provide the user with comprehensive SV annotations.

The field of genome-wide structural variation identification is rapidly advancing with LRS and OGM constantly evolving. However, currently both LRS and OGM technologies have limitations in identification of SNVs, large SVs >5kb (LRS) and small SVs <1kb (OGM) [17]. A combination of these various methods on the same genome will likely be necessary for optimal resolution and accuracy of SV detection [17, 47], which will require the design of integrated platforms able to detect and classify variants using multiple types of data sets. Similarly, accurate determination of the pathogenicity of SVs requires integration of multiple data sets (*e*.*g*. OGM and gene expression) as well as tools capable of annotating these various types of data sets. *nanotatoR* has this functionality and was able to considerably reduce analysis time compared to manual filtration of many steps. Future development of *nanotatoR* will be focused around adoption of variant annotation file (VCF) format as input files, support for SV calls produced by LRS/SRS technologies, additional population frequency databases such as gnomAD [48], and implementation of automated SV classification based on ACMG guidelines [49]. In addition, we plan to design a graphical interface for easy access and wider adoption.

## Availability and requirements

Project name: *nanotatoR*

Project home page: https://github.com/VilainLab/nanotatoR

Operating system(s): Platform-independent

Programming language: R (version >= 3.6)

Other requirements: Rtools (https://cran.r-project.org/bin/windows/Rtools/)

License: Modified BSD 3 (Berkley Software Distribution version 3).

Any restrictions to use by non-academics: license needed

## Declarations

### Ethics approval and consent to participate

Not Applicable.

### Consent for publication

Not Applicable.

### Availability of data and materials

The Bionano internal database was downloaded from using the following command wget http://bnxinstall.com/solve/Solve3.3_10252018.tar.gz. The DGV database was downloaded from http://dgv.tcag.ca/dgv/app/downloads?ref=GRCh37/hg19. The Decipher database was downloaded from https://decipher.sanger.ac.uk/files/downloads/population_cnv.txt.gz. The samples used for Examples I, II and building the internal database were downloaded from https://bionanogenomics.com/library/datasets/. The NA12878 RNAseq data was downloaded from https://www.ncbi.nlm.nih.gov/geo/query/acc.cgi?acc=GSM754335. Example III DMD data was used from [27].

## Acknowledgements

The authors are grateful to Vilain laboratory members for their valuable comments during the development of *nanotatoR*, especially Miguel Almalvez for coming up with the name *nanotatoR*.

## Authors’ contributions

SB and HB conceived the study. SB wrote the nanotatoR code. SB, HB, ED and EV provided key research concepts and wrote the manuscript. All the authors have read and approved the manuscript.

---

**Supplementary Table S1: NA24385 *nanotatoR*-annotated structural variant results** Content of the various tabs is detailed in the first tab (“LegendS1”) of the Excel workbook. SVs reported in the indel, inv, and trans tabs have undergone the default *nanotatoR* filtration (found in self molecules, passed chimeric score threshold). Columns in each sheet of the workbook are either a direct output of SVcaller or appended by nanotatoR, as indicated in column 3 of the table. Details about SVcaller columns are available at https://bionanogenomics.com/wp-content/uploads/2017/03/30041-SMAP-File-Format-Specification-Sheet.pdf

**Supplementary Table S2**: **Primary gene list generated for NA24385 (GM24385) dataset filtration using the term Osteoporosis**. A total of 307 genes were extracted (the *UGT2B17* gene is highlighted).

Columns show:

- **Genes**: Primary gene symbols.
- **Terms:** Query term associated with each gene. Each cell shows the gene symbol and, between parentheses, the query term and the database (Gene, OMIM, GTR or ClinVar) from which each gene was extracted.
- **Clinical Significance:** Only genes for which likely pathogenic or pathogenic variants were found in ClinVar are displayed in this column. The parentheses match the database order in column 2, displayed as (-, -, Pathogenic/Likely Pathogenic) to indicate the information is from ClinVar and not Gene or OMIM.

**Supplementary Table S3: Calculation of internal frequency for the deletion variant overlapping *UGT2B17* in sample NA24385**

Content of the tabs and column names is detailed in the first tab (“LegendS3”) of the Excel workbook. The columns used to calculate the filtered and unfiltered frequencies are highlighted in the del_totalData tab.

**Supplementary Table S4: Calculation of BNDB database frequency for the deletion variant overlapping *UGT2B17* in sample NA24385**

Content of the tabs and column names is detailed in the first tab (“LegendS4”) of the Excel workbook. The columns used to calculate the filtered and unfiltered frequencies are highlighted in the GM24385_data_all tab.

**Supplementary Table S5: Typical run times in minutes for the various tasks for the control sample datasets**. *nanotatoR* functions are indicated in italics in Column 1. Processing times for the Ashkenazi trio DLE labeling dataset (Example I) are shown in Column 1 (no RNA-Seq data was available for this sample). Processing times for the singleton sample (Example II) DLE labeling and dual enzyme labeling (SVmerge) are shown in columns 2 and 3 respectively.

**Supplementary Table S6: *nanotatoR* annotation of structural variants in the DLE-labeled sample NA12878 dataset**

Content of the various tabs is detailed in the first tab (“LegendS6”) of the Excel book. SVs reported in the indel, inv, and trans tabs have undergone the default *nanotatoR* filtration (found in self molecules, passed chimeric score threshold). Columns in each sheet of the workbook are either a direct output of SVcaller or appended by *nanotatoR*, as indicated in column 3 of the table. In the *all_PG_OV* tab, the cells containing the query genes are highlighted in purple. Details about SVcaller columns are available at https://bionanogenomics.com/wp-content/uploads/2017/03/30041-SMAP-File-Format-Specification-Sheet.pdf

**Supplementary Table S7: *nanotatoR* annotation of structural variants in the dual-labeled sample NA12878 dataset (SVmerge output)**

Content of the various tabs is detailed in the first tab (“LegendS7”) of the Excel book. SVs reported in the indel, inv, and trans tabs have undergone the default *nanotatoR* filtration (found in self molecules, passed chimeric score threshold). Columns in each sheet of the workbook are either a direct output of SVcaller or appended by *nanotatoR*, as indicated in column 3 of the table. In the *all_PG_OV* tab, the cells containing the query genes are highlighted in purple. Details about SVcaller columns are available at https://bionanogenomics.com/wp-content/uploads/2017/03/30041-SMAP-File-Format-Specification-Sheet.pdf

**Supplementary Table S8**: **Genes overlapping deletion variants for NA12878**

Chromosome and SV breakpoint start and end are shown in:

- NA12878 (Mak et.al. (2016) hg38 annotation): Overlapping genes coordinates identified in the original paper by Mak et al. 2016, with hg38 reference genome annotation.
- NA12878 (Mak et.al. (2016) hg19 *liftOver*): Overlapping genes coordinates identified in the original paper by Mak et al. 2016, lifted over to hg19.
- NA12878 (nanotatoR SVmerge annotation): Overlapping genes coordinates for NA12878 labelled by BSPQI and BSSSI enzyme (SVmerge), as annotated by *nanotatoR*.
- NA12878 (nanotatoR DLE annotation): Overlapping genes coordinates for NA12878 labelled by DLE1 enzyme (SVmerge), as annotated by *nanotatoR*.

**Supplementary Table S9: Variant annotation for DMD samples**

Singleton samples CDMD_1003 and CDMD_1159 were analyzed with single DLE labeling and are shown in the SingleLabel_Solo tab. Singleton samples CDMD_1155, CDMD_1156, and CDMD_1187 were analyzed with dual enzyme labeling and are shown in the DualLabel_Solo tab. Mother proband dyads CDMD_1131, CDMD_1157, and CDMD_1163 were analyzed with dual enzyme labeling and are shown in the DualLabelDuo tab. The columns used for Table 1 are highlighted.

Columns in each sheet of the workbook are either a direct output of SVcaller or appended by *nanotatoR*, as indicated in column 3 of the table in the first tab (“Legend S9”). Details about SVcaller columns are available at https://bionanogenomics.com/wp-content/uploads/2017/03/30041-SMAP-File-Format-Specification-Sheet.pdf

**Figure S1:**
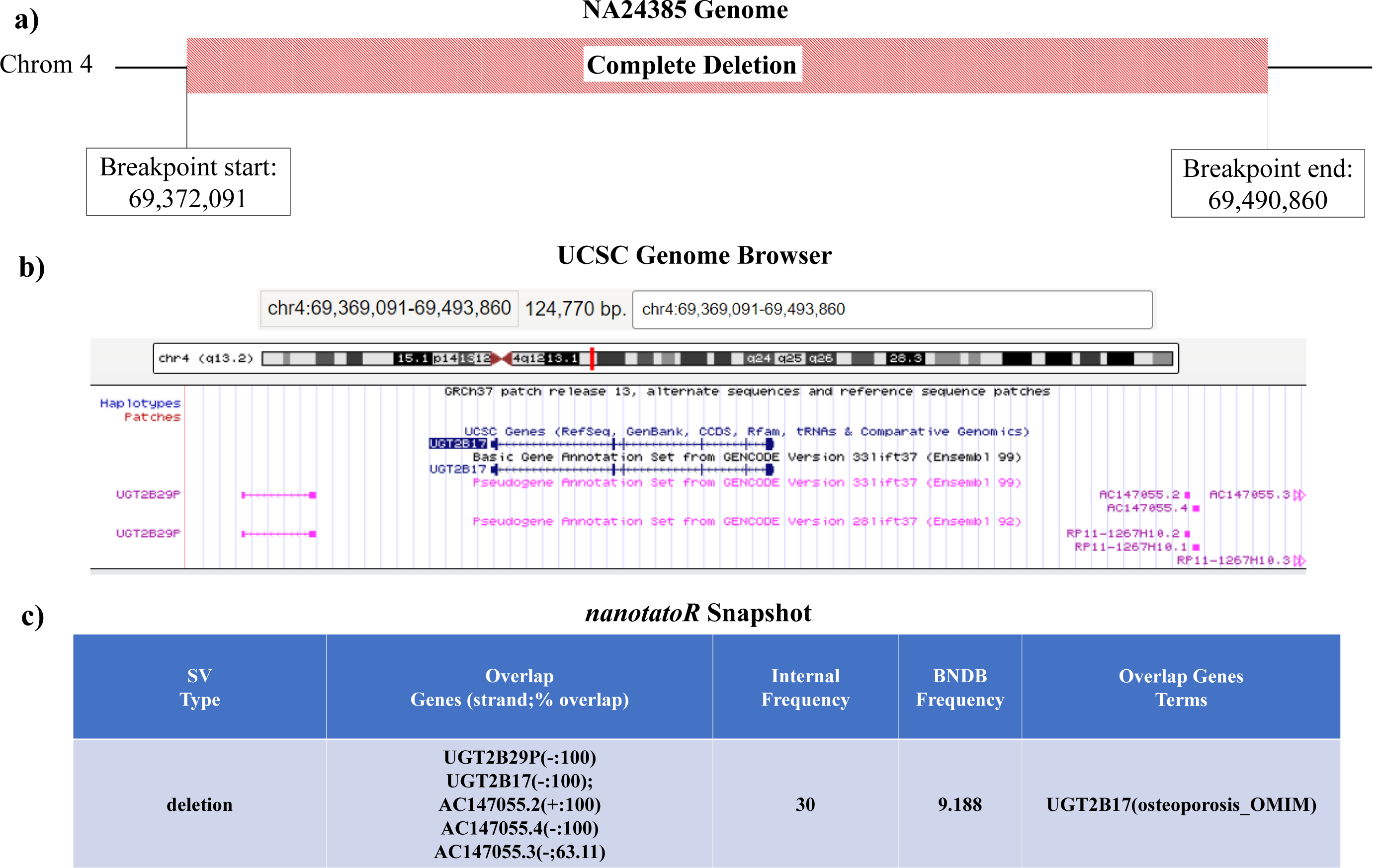
Deletion variant on chromosome 4 identified in sample NA24385: Cartoon of the chromosome 4 region deleted in the NA24385 genome **(a)** and screenshot of the matching UCSC genome browser output **(b)**. Breakpoints coordinates are shown as calculated by SVcaller in (**a**) and after including a method-average breakpoint error of +/−3kb [69,369,091 and 69,493,860] in (**b)**. This variant deletes the entirety of coding gene *UGT2B17*, as well as the 3 pseudogenes *UGT2B29P, AC147055*.*2* (or RP11-1267H10.2) and *AC147055*.*4* (or RP11-1267H10.1), and ∼63% of pseudogene *AC147055*.*3*. **c) *nanotatoR* snapshot:** The *nanotatoR* output indicates the overlap genes, the strand from which they are transcribed, and the percentage of the gene length overlapping with the deletion. It also displays the frequency (expressed as percentage) of the variant in the internal and BNDB databases, and the overlapping gene term with database info (osteoporosis found in the OMIM database).

## References

1. Goodwin S, McPherson JD, McCombie WR. Coming of age: ten years of next-generation sequencing technologies. Nat Rev Genet. 2016;17:333–51. doi: 10.1038/nrg.2016.49.

2. Bleidorn C. Third generation sequencing: technology and its potential impact on evolutionary biodiversity research. Syst Biodivers. 2016;14:1–8. doi: 10.1080/14772000.2015.1099575.

3. Lee H, Deignan JL, Dorrani N, Strom SP, Kantarci S, Quintero-Rivera F, et al. Clinical Exome Sequencing for Genetic Identification of Rare Mendelian Disorders. JAMA. 2014;312:1880. doi: 10.1001/jama.2014.14604.

4. Yang Y, Muzny DM, Reid JG, Bainbridge MN, Willis A, Ward PA, et al. Clinical Whole-Exome Sequencing for the Diagnosis of Mendelian Disorders. N Engl J Med. 2013;369:1502–11. doi: 10.1056/NEJMoa1306555.

5. Wright CF, FitzPatrick DR, Firth H V. Paediatric genomics: diagnosing rare disease in children. Nat Rev Genet. 2018;19:253–68. doi: 10.1038/nrg.2017.116.

6. Deelen P, van Dam S, Herkert JC, Karjalainen JM, Brugge H, Abbott KM, et al. Improving the diagnostic yield of exome-sequencing by predicting gene–phenotype associations using large-scale gene expression analysis. Nat Commun. 2019;10:2837. doi: 10.1038/s41467-019-10649-4.

7. Meienberg J, Bruggmann R, Oexle K, Matyas G. Clinical sequencing: is WGS the better WES? Hum Genet. 2016;135:359. doi: 10.1007/S00439-015-1631-9.

8. Belkadi A, Bolze A, Itan Y, Cobat A, Vincent QB, Antipenko A, et al. Whole-genome sequencing is more powerful than whole-exome sequencing for detecting exome variants. Proc Natl Acad Sci U S A. 2015;112:5473–8. doi: 10.1073/pnas.1418631112.

9. Kosugi S, Momozawa Y, Liu X, Terao C, Kubo M, Kamatani Y. Comprehensive evaluation of structural variation detection algorithms for whole genome sequencing. Genome Biol. 2019;20:117. doi: 10.1186/s13059-019-1720-5.

10. Guan P, Sung W-K. Structural variation detection using next-generation sequencing data: A comparative technical review. Methods. 2016;102:36–49. doi: 10.1016/j.ymeth.2016.01.020.

11. Cameron DL, Di Stefano L, Papenfuss AT. Comprehensive evaluation and characterisation of short read general-purpose structural variant calling software. Nat Commun. 2019;10:3240. doi: 10.1038/s41467-019-11146-4.

12. Amarillo IE, Nievera I, Hagan A, Huchthagowder V, Heeley J, Hollander A, et al. Integrated small copy number variations and epigenome maps of disorders of sex development. Hum genome Var. 2016;3:16012. doi: 10.1038/hgv.2016.12.

13. Jain M, Koren S, Miga KH, Quick J, Rand AC, Sasani TA, et al. Nanopore sequencing and assembly of a human genome with ultra-long reads. Nat Biotechnol. 2018;36:338–45. doi: 10.1038/nbt.4060.

14. Cretu Stancu M, van Roosmalen MJ, Renkens I, Nieboer MM, Middelkamp S, de Ligt J, et al. Mapping and phasing of structural variation in patient genomes using nanopore sequencing. Nat Commun. 2017;8:1326. doi: 10.1038/s41467-017-01343-4.

15. Ebbert MTW, Jensen TD, Jansen-West K, Sens JP, Reddy JS, Ridge PG, et al. Systematic analysis of dark and camouflaged genes reveals disease-relevant genes hiding in plain sight. Genome Biol. 2019;20:97. doi: 10.1186/s13059-019-1707-2.

16. Zhou A, Lin T, Xing J. Evaluating nanopore sequencing data processing pipelines for structural variation identification. Genome Biol. 2019;20:237. doi: 10.1186/s13059-019-1858-1.

17. Chaisson MJP, Sanders AD, Zhao X, Malhotra A, Porubsky D, Rausch T, et al. Multiplatform discovery of haplotype-resolved structural variation in human genomes. Nat Commun. 2019;10:1784. doi: 10.1038/s41467-018-08148-z.

18. Levy-Sakin M, Ebenstein Y. Beyond sequencing: optical mapping of DNA in the age of nanotechnology and nanoscopy. Curr Opin Biotechnol. 2013;24:690–8. doi: 10.1016/j.copbio.2013.01.009.

19. Levy-Sakin M, Pastor S, Mostovoy Y, Li L, Leung AKY, McCaffrey J, et al. Genome maps across 26 human populations reveal population-specific patterns of structural variation. Nat Commun. 2019;10:1025. doi: 10.1038/s41467-019-08992-7.

20. Hastie AR, Lam ET, Pang AWC, Zhang LX, Andrews W, Lee J, et al. Rapid Automated Large Structural Variation Detection in a Diploid Genome by NanoChannel Based Next-Generation Mapping. bioRxiv. 2017;:102764. doi: 10.1101/102764.

21. Bocklandt S, Hastie A, Cao H. Bionano Genome Mapping: High-Throughput, Ultra-Long Molecule Genome Analysis System for Precision Genome Assembly and Haploid-Resolved Structural Variation Discovery. Adv Exp Med Biol. 2019;1129:97–118. doi: 10.1007/978-981-13-6037-4_7.

22. Leung AK-Y, Kwok T-P, Wan R, Xiao M, Kwok P-Y, Yip KY, et al. OMBlast: alignment tool for optical mapping using a seed-and-extend approach. Bioinformatics. 2016;33:btw620. doi: 10.1093/bioinformatics/btw620.

23. Leung AK-Y, Jin N, Yip KY, Chan T-F. OMTools: a software package for visualizing and processing optical mapping data. Bioinformatics. 2017;33:2933–5. doi: 10.1093/bioinformatics/btx317.

24. Jaratlerdsiri W, Chan EKF, Petersen DC, Yang C, Croucher PI, Bornman MSR, et al. Next generation mapping reveals novel large genomic rearrangements in prostate cancer. Oncotarget. 2017;8:23588–602. doi: 10.18632/oncotarget.15802.

25. Du C, Mark D, Wappenschmidt B, Böckmann B, Pabst B, Chan S, et al. A tandem duplication of BRCA1 exons 1–19 through DHX8 exon 2 in four families with hereditary breast and ovarian cancer syndrome. Breast Cancer Res Treat. 2018;172:561–9. doi: 10.1007/s10549-018-4957-x.

26. Dixon JR, Xu J, Dileep V, Zhan Y, Song F, Le VT, et al. Integrative detection and analysis of structural variation in cancer genomes. Nat Genet. 2018;50:1388–98. doi: 10.1038/s41588-018-0195-8.

27. Barseghyan H, Tang W, Wang RT, Almalvez M, Segura E, Bramble MS, et al. Next-generation mapping: a novel approach for detection of pathogenic structural variants with a potential utility in clinical diagnosis. Genome Med. 2017;9:90. doi: 10.1186/s13073-017-0479-0.

28. Dai Y, Li P, Wang Z, Liang F, Yang F, Fang L, et al. Single-molecule optical mapping enables accurate molecular diagnosis of facioscapulohumeral muscular dystrophy (FSHD). bioRxiv. 2018;:286104. doi: 10.1101/286104.

29. Sharim H, Grunwald A, Gabrieli T, Michaeli Y, Margalit S, Torchinsky D, et al. Long-read single-molecule maps of the functional methylome. Genome Res. 2019;29:646–56. doi: 10.1101/gr.240739.118.

30. Dobin A, Davis CA, Schlesinger F, Drenkow J, Zaleski C, Jha S, et al. STAR: ultrafast universal RNA-seq aligner. Bioinformatics. 2013;29:15–21. doi: 10.1093/bioinformatics/bts635.

31. Li B, Dewey CN. RSEM: accurate transcript quantification from RNA-Seq data with or without a reference genome. BMC Bioinformatics. 2011;12:323. doi: 10.1186/1471-2105-12-323.

32. MacDonald JR, Ziman R, Yuen RKC, Feuk L, Scherer SW. The Database of Genomic Variants: a curated collection of structural variation in the human genome. Nucleic Acids Res. 2014;42 Database issue:D986–92. doi: 10.1093/nar/gkt958.

33. Firth H V, Richards SM, Bevan AP, Clayton S, Corpas M, Rajan D, et al. DECIPHER: Database of Chromosomal Imbalance and Phenotype in Humans Using Ensembl Resources. Am J Hum Genet. 2009;84:524–33. doi: 10.1016/j.ajhg.2009.03.010.

34. Bionano Genomics. Bionano Solve Theory of Operation: Variant Annotation Pipeline. 2018. https://bionanogenomics.com/wp-content/uploads/2018/04/30190-Bionano-Solve-Theory-of-Operation-Variant-Annotation-Pipeline.pdf. Accessed 19 Feb 2020.

35. Landrum MJ, Lee JM, Benson M, Brown GR, Chao C, Chitipiralla S, et al. ClinVar: improving access to variant interpretations and supporting evidence. Nucleic Acids Res. 2018;46:D1062–7. doi: 10.1093/nar/gkx1153.

36. Rubinstein WS, Maglott DR, Lee JM, Kattman BL, Malheiro AJ, Ovetsky M, et al. The NIH genetic testing registry: a new, centralized database of genetic tests to enable access to comprehensive information and improve transparency. Nucleic Acids Res. 2012;41:D925–35. doi: 10.1093/nar/gks1173.

37. David J. Winter. rentrez: an R package for the NCBI eUtils API. R J. 2017;9:520--526. https://cran.r-project.org/web/packages/rentrez/citation.html. Accessed 1 Aug 2019.

38. Cao Z, Wang L, Chen Y, Cai R, Lu J, Yu Y, et al. VarfromPDB: An Automated and Integrated Tool to Mine Disease-Gene-Variant Relations from the Public Databases and Literature. J Proteomics Bioinform. 2017;10:311–5. doi: 10.4172/jpb.1000455.

39. Walker A. openxlsx: Read, Write and Edit XLSX Files. R package version 4.1.0. https://CRAN.R-project.org/package=openxlsx. 2018;:2018. https://cran.r-project.org/web/packages/openxlsx/index.html. Accessed 5 Aug 2019.

40. Bionano Genomics. SMAP File Format Specification Sheet. 2019. https://bionanogenomics.com/wp-content/uploads/2017/03/30041-SMAP-File-Format-Specification-Sheet.pdf. Accessed 25 Feb 2020.

41. Hastie A, Liang T, Pham K, Saghbini M, Džakula Ž, Cao H. De Novo Assembly of the Genome-in-a-Bottle Reference Ashkenazi Trio, Structural Variation Discovery and Comparison with Other Individuals by Next-Generation Mapping. In: ASHG. 2015. https://bionanogenomics.com/wp-content/uploads/2017/11/Bionano-Poster_ASHG2015_Alex_De-Novo-Assembly-Genome-in-a-Bottle-Reference-Ashkenazi-Trio.pdf. Accessed 24 Feb 2020.

42. Yang T-L, Chen X-D, Guo Y, Lei S-F, Wang J-T, Zhou Q, et al. Genome-wide copy-number-variation study identified a susceptibility gene, UGT2B17, for osteoporosis. Am J Hum Genet. 2008;83:663–74. doi: 10.1016/j.ajhg.2008.10.006.

43. Mak ACY, Lai YYY, Lam ET, Kwok T-P, Leung AKY, Poon A, et al. Genome-Wide Structural Variation Detection by Genome Mapping on Nanochannel Arrays. Genetics. 2016;202:351–62. doi: 10.1534/genetics.115.183483.

44. Weischenfeldt J, Symmons O, Spitz F, Korbel JO. Phenotypic impact of genomic structural variation: insights from and for human disease. Nat Rev Genet. 2013;14:125–38. doi: 10.1038/nrg3373.

45. Geoffroy V, Herenger Y, Kress A, Stoetzel C, Piton A, Dollfus H, et al. AnnotSV: an integrated tool for structural variations annotation. Bioinformatics. 2018;34:3572–4. doi: 10.1093/bioinformatics/bty304.

46. Wang K, Li M, Hakonarson H. ANNOVAR: functional annotation of genetic variants from high-throughput sequencing data. Nucleic Acids Res. 2010;38:e164–e164. doi: 10.1093/nar/gkq603.

47. Brandler WM, Antaki D, Gujral M, Kleiber ML, Whitney J, Maile MS, et al. Paternally inherited cis-regulatory structural variants are associated with autism. Science. 2018;360:327–31. doi: 10.1126/science.aan2261.

48. Carlson M. org.Hs.eg.db: Genome wide annotation for Human. 2019. https://bioconductor.org/packages/release/data/annotation/html/org.Hs.eg.db.html. Accessed 1 Aug 2019.

49. Richards S, Aziz N, Bale S, Bick D, Das S, Gastier-Foster J, et al. Standards and guidelines for the interpretation of sequence variants: a joint consensus recommendation of the American College of Medical Genetics and Genomics and the Association for Molecular Pathology. Genet Med. 2015;17:405–24. doi: 10.1038/gim.2015.30.

